# Defining the *in vivo* role of mTORC1 in thyrocytes by studying the TSC2 conditional knockout mouse model

**DOI:** 10.1101/2023.11.01.565171

**Authors:** Camila Rossetti, Bruna Lourençoni, Flavia Peçanha, Aime T Franco, Vania Nosé, Everardo Carneiro, John Lew, Ernesto Bernal-Mizrachi, Joao Pedro Werneck-de Castro

## Abstract

The thyroid gland is susceptible to abnormal epithelial cell growth, often resulting in thyroid dysfunction. The serine-threonine protein kinase mechanistic target of rapamycin (mTOR) regulates cellular metabolism, proliferation, and growth through two different protein complexes, mTORC1 and mTORC2. The PI3K-Akt-mTORC1 pathway’s overactivity is well associated with heightened aggressiveness in thyroid cancer, but recent studies indicate the involvement of mTORC2 as well. To elucidate mTORC1’s role in thyrocytes, we developed a novel mouse model with mTORC1 gain of function in thyrocytes by deleting Tuberous Sclerosis Complex 2 (TSC2), an intracellular inhibitor of mTORC1. The resulting *TPO-TSC2^KO^* mice exhibited a significant reduction in TSC2 levels, leading to a six-fold increase in mTORC1 activity. Thyroid glands of both male and female *TPO-TSC2^KO^* mice displayed rapid enlargement and continued growth throughout life, accompanied by heterogeneity among thyroid follicles, larger follicles, increased colloid and epithelium. We observed elevated thyrocyte proliferation as indicated by Ki67 staining and elevated Cyclin D3 expression in the *TPO-TSC2^KO^* mice. mTORC1 activation resulted in a progressive downregulation of key genes involved in thyroid hormone (TH) biosynthesis, including thyroglobulin, thyroid peroxidase, and sodium-iodide symporter (NIS), while TTF1, PAX8, and MCT8 mRNA levels remained unaffected. NIS protein expression was also diminished in *TPO-TSC2^KO^* mice. Treatment with the mTORC1 inhibitor rapamycin prevented thyroid mass expansion and restored the gene expression alterations in *TPO-TSC2^KO^* mice. Although T_4_, T_3_ and TSH plasma levels were normal at 2 months of age, a slight decrease in T_4_ and an increase in TSH levels were observed at 6 and 12 months of age while T3 remained similar in *TPO-TSC2^KO^* compared to littermate control mice. *TPO-TSC2^KO^* mice aged to 12 months or older developed aberrant thyroid conditions, including follicular hyperplasia, inflammation, and thyroid tumors. In conclusion, our thyrocyte-specific mouse model reveals that mTORC1 activation inhibits TH biosynthesis, suppresses thyrocyte gene expression, and promotes growth and proliferation. Chronic mTORC1 activation leads to thyroid tumor formation, highlighting the role of mTORC1 in thyroid dysfunction and tumorigenesis.

## INTRODUCTION

The thyroid gland is often the site of abnormal growth and proliferation of epithelial cells, which can result in thyroid dysfunction [2-5]. A total of $2.1-4.3 billion was spent on treatment for thyroid dysfunction among adult women [6]. Cost-effectiveness analyses have estimated that the screening and management of all thyroid nodules in the United States would incur $25.1 billion in costs [7]. For nodules representing malignant disease, the costs of well-differentiated thyroid cancer care in the United States are projected to exceed $3.5 billion by 2030 [5, 7].

The serine–threonine protein kinase mechanistic target of rapamycin (mTOR) plays a crucial role in regulating cellular metabolism, growth, and proliferation [8]. mTOR forms two distinct multi-protein complexes: mTOR complex 1 (mTORC1) and mTOR complex 2 (mTORC2). mTORC1 comprises mTOR, Raptor, PRASA40, mLST8, and Deptor, while mTORC2 includes mTOR, Rictor, mSIN1, mLST8, and Deptor. These complexes have unique targets and signaling roles in the cell [9]. mTORC1, acting as a downstream effector for growth factors, hormones, cellular energy, and nutrients—particularly amino acids—integrates information on nutritional abundance and environmental status to balance anabolism and catabolism [10-14]. Conversely, mTORC2 governs cytoskeletal behavior and activates pro-survival pathways [8, 15, 16]. The mTORC1 pathway’s regulation involves the tuberous sclerosis complex (TSC1/2), serving as a GTPase-activating protein for Rheb. This complex catalyzes the conversion from the active Rheb-GTP state to the inactive GDP-bound state [17-19]. Activation of mTORC1 requires binding to active Rheb-GTP, with TSC1/2 acting as a critical’molecular brake’ [9]. mTORC1 tightly regulates protein synthesis by phosphorylating eukaryotic initiation factor 4E-binding proteins (4E-BPs) and p70S6 kinase 1 (S6K). Phosphorylation of 4E-BP1 enhances 5′ cap-dependent translation of mRNA [20, 21], while activation of S6K promotes ribosomal synthesis and accelerates translation elongation [22, 23]. By regulating all necessary cellular metabolic pathways, mTORC1 is essential for cell growth and supports proliferation.

Tuberous Sclerosis (TSC) is an autosomal dominant disorder resulting from mutations in TSC1 or TSC2, leading to hyperactivation of the mTORC1 pathway [24, 25]. Dysregulated mTORC1 signaling in these patients leads in increased cell growth and proliferation, manifesting as benign tumors in the brain, skin, heart, lungs, and kidneys [26]. Neurological symptoms, including seizures, autism spectrum disorder, and cognitive disability, are also observed [27]. The incidence of TSC is approximately 1 case per 6,000–10,000 live births [28]. Emerging evidence suggests that the thyroid gland is a target of TSC [29-35]. In routine chest computed tomography for lung disease assessment, 20% of TSC patients exhibited thyroid nodules, with half of them diagnosed with multiple nodules [30]. Thyroid abnormalities, although less frequent, in TSC encompass papillary and medullary thyroid carcinoma [29, 31], as well as hypothyroidism [32, 33].

Thyroid Stimulating Hormone (TSH) released by the pituitary gland regulates virtually all thyroid biological functions, including steps in thyroid hormone (TH) synthesis, secretion, and thyrocyte growth and proliferation [36-38]. TSH also impacts non-epithelium cells such as endothelial and pericytes, promoting vasculature function and expansion [37]. TSH effects on thyrocytes are primarily mediated through activation of adenylate cyclase by the alpha-subunit of stimulatory GTP-binding protein (Gαs), but it also activates the Gαq/phospholipase C cascade [39-42]. Specifically, TSH stimulates the expression of several key components involved in TH production: the sodium iodide symporter (NIS), thyroid peroxidase (TPO), thyroglobulin (TG), and it aids in the generation of H_2_O_2_ [43-46]. TSH increases the T_4_-to-T_3_ conversion and accelerates the internalization of colloid-containing TG pinocytic vesicles by thyrocytes [47-50].

In addition to enhancing thyrocyte function, TSH promotes thyrocyte proliferation and growth [37], partly mediated by activation of the mTORC1 pathway [51-54]. Notably, *in vitro* studies have shown that the TSH receptor exhibits intrinsic phosphatidyl-inositol-3-kinase (PI3K) activity, activating mTORC1 to promote thyrocyte proliferation [55]. Furthermore, TSH induces mTORC1 activation by PKA-mediated phosphorylation of PRAS40 [54]. mTORC1 is also activated by Insulin (INS) and Insulin-like Growth Factor (IGF1) through the PI3K-AKT-TSC2-Rheb-mTORC1 axis [54, 55], cooperating with TSH to promote thyrocyte proliferation [56-58]. *In vivo*, mTORC1 activity is enhanced in the thyroid of mice undergoing diet-induced goiter (high endogenous circulating TSH levels) [53-55, 59]. Its systemic inhibition by rapamycin reduces thyrocyte proliferation but not growth [53] and decreases follicle activity in mice [55]. Data from transgenic mouse models also indirectly suggest that mTORC1 could be the hub for TSH and INS/IGF1 signals, regulating thyrocyte proliferation and growth. Thyrocyte-specific IGF1 receptor genetic ablation (*IGF1R^KO^*) impairs iodine deficiency (ID)- induced mTORC1 activation, hindering thyroid expansion [59]. This suggests that TSH relies on the IGF1 signaling pathway to fully activate mTORC1 during adaptive responses to ID. However, contradictory reports indicate that *IGF1R^KO^* mice [60] and IGF1 and INS receptor double knockout mice (*IGF1RKO/IR^KO^*) [61] have intense thyrocyte proliferation and growth, and mTORC1 activation [61]. The authors suggested that TSH compensates for the lack of INS/IGF1 signaling [60, 61]. Although indirect, these data suggest mTORC1 could be playing a role in the regulation of thyrocyte proliferation and growth in response to TSH and INS/IGF1. However, the direct *in vivo* effects of mTORC1 on thyrocytes was not addressed so far.

While mTORC1 pathway has been studied in the context of thyrocyte proliferation and growth, its role in TH synthesis and thyroid function remains unclear [62-64].The regulation of NIS expression and activity exemplifies this controversy. Studies demonstrated that IGF1 inhibits NIS expression in rat FRTL-5 and PCCL3 thyrocytes [42, 52, 65]. This inhibitory effect has been shown to be mediated by PI3K [42, 52, 65] and mTORC1 [65]. However, recent studies in primary human thyrocytes show that TSH and IGF1 elicit additive effects on TPO, TG, and the type-1 deiodinase (Dio1) expression, with synergistic effects on NIS mRNA and protein [63, 64]. Whether mTORC1 controls NIS expression and thyrocyte differentiation is clinically relevant because the loss of NIS expression in thyroid cancer is the proposed mechanism for the refractory response to radioiodine therapy [66]. Moreover, several common oncogenic signaling pathways converge on mTORC1 [9, 67, 68] and evidence indicates its involvement in the progression of more lethal forms of thyroid cancer [69-74]. However, recent data in papillary thyroid carcinoma indicate that mTORC2, but not mTORC1, is overactivated and suppresses NIS expression [70, 71].

The discrepancy in these findings likely stems from the lack of *in vivo* animal models to study mTORC1 or mTORC2 induction. Our current understanding of mTORC1’s role in thyrocytes is primarily based on *in vitro* studies, which do not fully recapitulate *in vivo* TH synthesis and secretion due to the disruption of the critical angio-follicular unit and cell polarity [37]. Additionally, cultured thyroid cancer cell lines are not only of thyroid origin and lose normal functions such as TSH responsiveness [75, 76]. To assess the impact of mTORC1 activation on thyrocyte gene expression, we generated a mouse model of mTORC1 gain of function by specifically deleting the TSC2 gene in thyrocytes (*TPO-TSC2^KO^* mice). These mice exhibit highly active mTORC1 and low mTORC2 activity in the thyroid gland, allowing us to directly investigate mTORC1 biological effects. mTORC1 activation led to thyrocyte growth and proliferation independent of circulating TSH. The thyroids of the *TPO-TSC2^KO^* mice had decreased expression of essential genes for TH biosynthesis (*NIS, TPO*, and *TG*) but normal TTF1, PAX8 and MTC8 expression. Treatment with rapamycin in these mice confirmed the mTORC1-targeted genes in thyrocytes. Aged *TPO-TSC2^KO^* mice (12 months or older) presented thyroid tumors. Collectively, these data establish mTORC1 as a crucial intracellular component implicated in TH biosynthesis.

## MATERIAL AND METHODS

### Animals

All experimental procedures were planned following the American Thyroid Association guide to investigating TH economy and action in rodent and cell models [77] and approved by the local Institutional Animal Care and Use Committee (IACUC 20-166). All mice were maintained at 22 ± 1 °C on a 12h light–dark cycle, with free access to food and water. Thyroid-specific *TSC2* knockout mice (*TPO-TSC2^KO^*) were generated by crossing *Tsc2*^*flox/flox*^ mice with mice expressing the Cre recombinase gene under the control of the TPO promoter. The *Tsc2*^*flox/flox*^ mice harbor a modified endogenous *TSC2* gene with *loxP* sites flanking exons 2 to 4 [1] (Fig 1A). TPO-cre mouse was kindly donated by Dr. Shioko Kimura [78]. Experiments were performed on male and female mice that were on a mixed 129 x C57BL/6 background. We used littermates *Tsc2*^*flox/flox*^ *or TPO-cre* mice as controls. Rapamycin-treated mice received daily intraperitoneal injections of either 2 mg/Kg of b.w. Rapa (Cayman Chemical Company, Ann Harbor, MI) or vehicle (5.2% Poly(ethylene glycol) and 5.2% Tween 80; Sigma-Aldrich, St. Louis, MO) from weaning (3 weeks of age) to 2 months of age.

**Figure 1.**
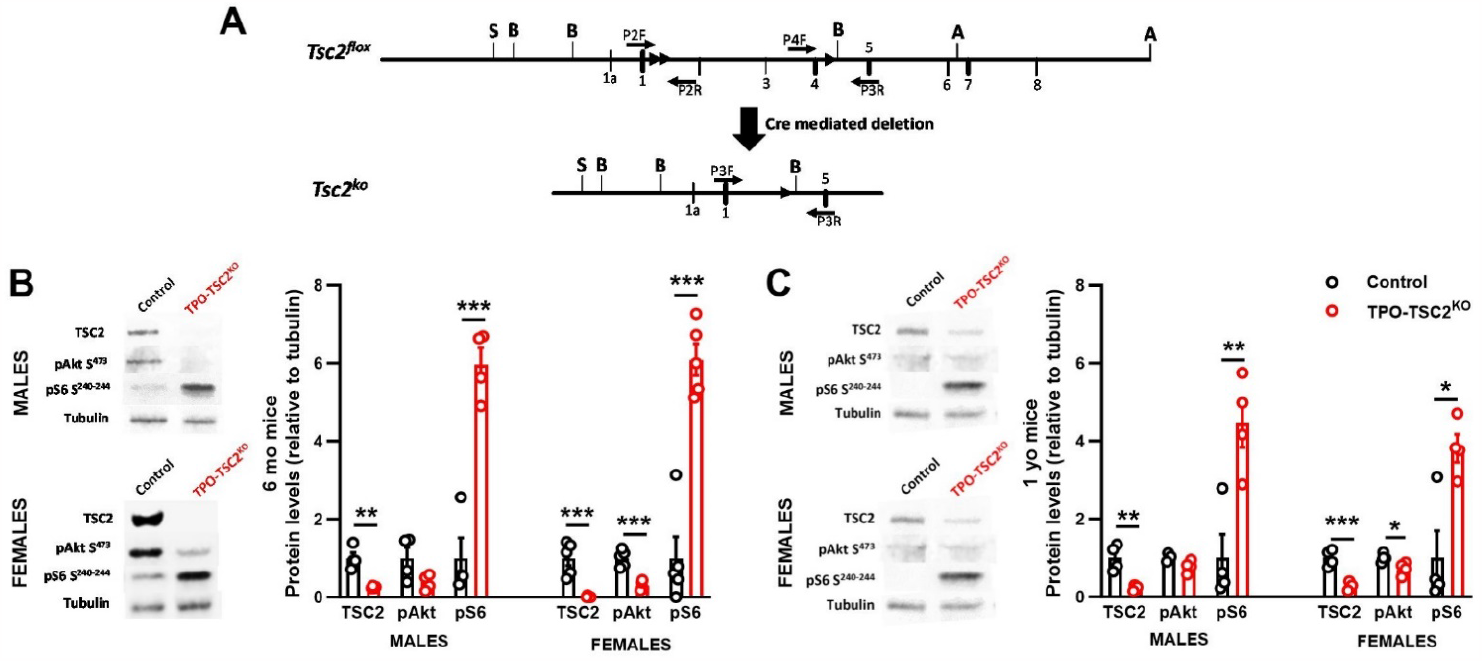
The TPO-TSC2 knockout model. **(a)** Gene-targeting construct from Hernandez, 2007 [1]. Exons are indicated by black boxes labeled with exon numbers. LoxP sites are marked by black triangles. The loxP-frt-neo-frt cassette was inserted in intron 1 and a loxP-BamHI site in intron 4. Cre-mediated recombination excises 2, 3 and 4; **(b)** Western blotting of the thyroid gland of 6 months old males and females *TPO-TSC2^KO^* and littermate mice for TSC2, pAkt Ser^473^ (mTORC2 activity) and pS6 Ser^240-244^ (mTORC1 activity) protein expression and **(c)** Same as in **b** except that thyroid of 1-year old mice were assessed. N = 4-5 mice per group. Data are shown as mean ± SEM. * p < 0.05; ** p < 0.01; *** p < 0.0001 compared to control mice determined by unpaired two-tail Student’s t-test.

### Body composition and hormone measurements

Body weight and glycemia were followed monthly from weaning to 12 months of age. Dual-energy X-ray absorptiometry (DEXA; Lunar Pixi, Janesville, WI) determined lean body mass and fat mass. For that, mice were fasted for 4-5 hours and anesthetized with ketamine-xylazine (36 mg/kg and 4 mg/kg) before imaging. Blood samples were collected in K_2_EDTA-coated tubes and centrifuged at 1,000 x g for 10 minutes within 30 minutes of collection. Total Thyroxine (T_4_) and total Triiodothyronine (T_3_) were measured using 25 µL of plasma and ELISA kits from Cusabio (Houston, TX). TSH levels were determined using Luminex multiplex technologies with one-plex beads (Milliplex Mouse Pituitary Magnetic Bead Panel, Millipore, Burlington, MA). All hormone determinations were performed according to the manufacturer’s instructions.

### Western blotting

One thyroid lobe per animal was lysed in buffer containing 125 mM Tris (pH 6.8), 1 mM dithiothreitol, 2% SDS, 1 µM phenylmethylsulfonyl fluoride, and protease and phosphatase inhibitor cocktails (Roche Diagnostics, Indianapolis, IN). Protein quantity was measured using the bicinchoninic acid (BCA) assay method and 30 µg of protein were separated by electrophoresis into 4-20% gradient SDS–PAGE gels. Then, proteins were transferred to polyvinylidene difluoride (PVDF) membranes (Millipore, Burlington, MA) overnight. After 1 h in blocking buffer, membranes were incubated overnight at 4 °C with primary antibodies listed in Table 1. Immunocomplexes were detected using fluorescent secondary antibodies and the Odissey FC Imager (LI-COR Biosciences, Lincoln, NE). Bands were submitted to densitometric analysis by ImageJ 1.43u software and normalized to tubulin, β-actin or total S6 (tS6). Images were cropped for presentation.

**TABLE 1.** Antibodies used for western blotting.

| Antibody | Specie | Source | Catalog | Dilution |
| --- | --- | --- | --- | --- |
| p-Akt Ser<sup>473</sup> | Rabbit | Cell Signaling | #4060 | 1:1000 |
| β-Actin | Mouse | Sigma-Aldrich | #A5441 | 1:5000 |
| Cyclin D2 | Rabbit | Cell Signaling | #3741S | 1:1000 |
| Cyclin D3 | Mouse | Cell Signaling | #2936S | 1:1000 |
| NIS | Rabbit | Millipore | #ABC1453 | 1:1000 |
| p-S6 Ser<sup>240-244</sup> | Rabbit | Cell Signaling | #5364 | 1:5000 |
| Total S6 | Mouse | Cell Signaling | #2317S | 1:750 |
| Tuberin/TSC2 | Rabbit | Cell Signaling | #4308 | 1:1000 |
| Tubulin | Mouse | Sigma-Aldrich | #T5168 | 1:5000 |

### Real-time PCR

One thyroid lobe was used for total RNA extraction according to the manufacturer’s protocol (RNeasy Micro Kit, Qiagen, Hilden, Germany). RNA concentration and purity were determined by spectrophotometry (ratio 260/280 nm). cDNA was synthesized using a reverse transcriptase reaction (High-Capacity cDNA Reverse Transcription kit, Applied Biosystems, Waltham, MA). Real-time PCRs were run in duplicate using PerfeCTa SYBR Green FastMix ROX gene expression assay (Quantabio, Beverly, MA) in a QuantStudio 3 real-time PCR system (Applied Biosystems, Waltham, MA). mRNA expression was calculated using the relative standard curve method and normalized for the housekeeping gene *18S* or *CycloB.* Primers were validated using a standard curve prepared with a mixture of cDNA from all samples (r^2^ > 0.98 and efficiency ∼80-110%) and are listed in Table 2.

**TABLE 2.**
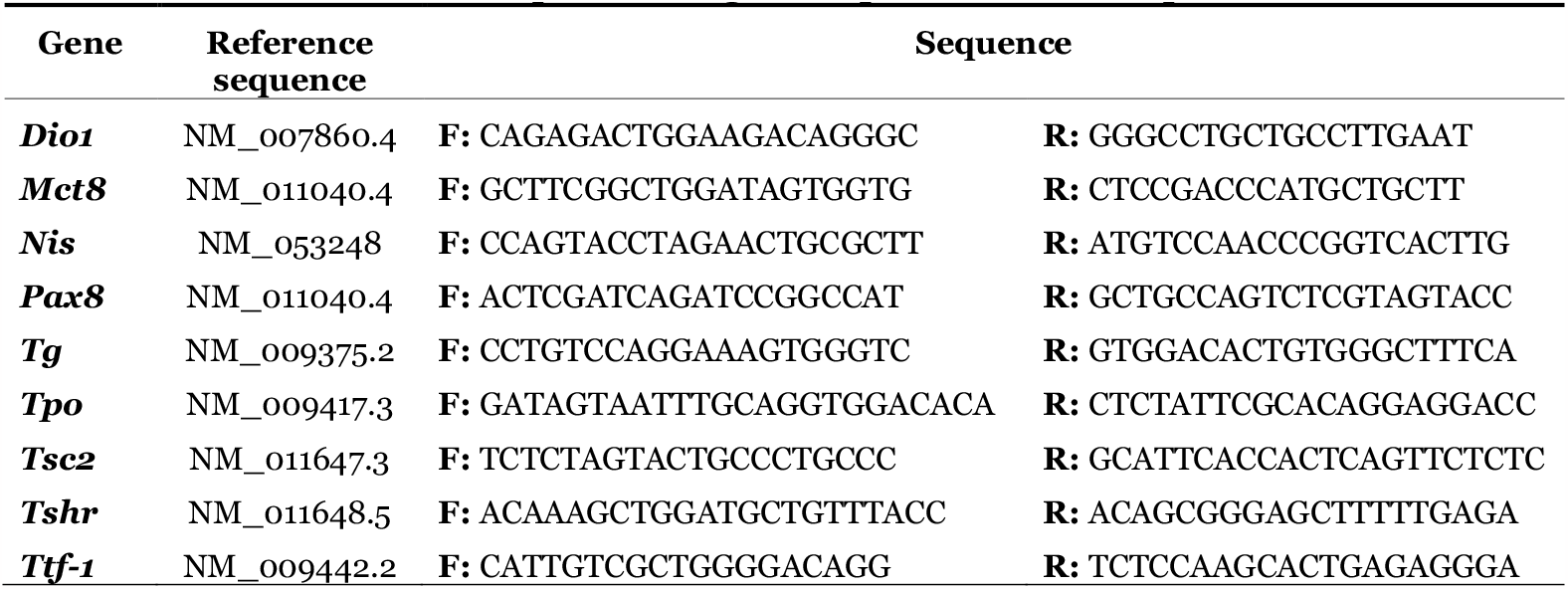
Primer sequences for gene amplification via RT-qPCR.

### Histological analysis

Thyroids were collected and fixed in 4% paraformaldehyde overnight, followed by dehydration in ethanol. Tissues were embedded in paraffin and 5 µm sections were stained with Periodic Acid-Schiff (PAS) or Hematoxylin and Eosin for morphometric analysis, according to [77, 79, 80]. High-resolution images were obtained using a digital camera (DFC360FX camera, Leica Microsystems, Wetzlar, Germany) coupled to a microscope (Leica DM5500B, Leica Microsystems). Images were captured with Leica Application Suite X (LAS X). PAS images were used for the quantification of follicular, colloid, and epithelium (total area – colloid area) areas [77, 79, 80]. At least 100 follicles per mouse were included in the analysis. All the measurements were performed using the ImageJ software.

### Thyrocyte proliferation quantification

Mouse thyroids attached to the trachea were fixed by immersion in 4% paraformaldehyde overnight at 4° and embedded in paraffin. To determine the number of proliferating thyrocytes, 5 µm sections were stained for Ki-67 and TTF-1 (nuclear thyroid marker) using anti-Mo/Rt Ki-67 (Invitrogen, Waltham, MA) and anti-TTF-1 (D2E8) Rabbit mAb (Cell Signaling Technology, Danvers, MA) primary antibodies. Fluorescent images were captured with a Leica DM5500B microscope equipped with a motorized stage and a DFC360FX camera (Leica Microsystems, Wetzlar, Germany). The microscope was controlled by the OASIS-blue PCI controller and the Leica Application Suite X (LAS X). Ki-67^+^ and TFF-1^+^ cells were quantified by ImageJ.

### Statistical analysis

For normal distributed data, results are presented as mean ± standard error of the mean (SEM) and analyzed as follows: 1) Student’s t-test for comparisons between 2 groups or 2) *two-way* ANOVA to analyze the effect of two independent factors followed by Tukey post-hoc analysis. Non-normal distributed data are presented as median and quartiles (Violin plot), and the Mann-Whitney U test was used to compare two groups. Results were considered statistically significant if p < 0.05. Statistical analyses and graphing were performed using GraphPad Prism version 9.4 (GraphPad Software, San Diego, CA, US).

## RESULTS

### Activation of mTORC1 in thyrocytes promotes thyroid gland growth

To investigate the role of mTORC1 pathway in thyrocytes, we generated a new mouse model of mTORC1 gain of function specifically in thyrocytes by crossing the TSC2-floxed mouse with the TPO-cre mouse (*TPO-TSC2^KO^* mice; Fig 1A). Thyroids of *TPO-TSC2^KO^* mice had a 70-80% decrease in TSC2 protein levels and a 6-fold induction of mTORC1 activity (pS6) in both females and males (Fig 1). In contrast, mTORC2 activity decreased in the *TPO-TSC2^KO^* mice thyroids (lower pAkt Ser^473^ phosphorylation). Male and female *TPO-TSC2^KO^* mice did not gain weight as littermate control mice and had less fat mass at 12 months of age (Fig 2A-C). mTORC1 overactivation promoted a remarkable thyroid gland enlargement as early as 1 month of age (Fig 2D). Thyroid mass was higher in TSC2 knockout mice compared to the thyroid of control animals at 1, 2, 6, and 12 months of age (Fig 2D-F).

**Figure 2.**
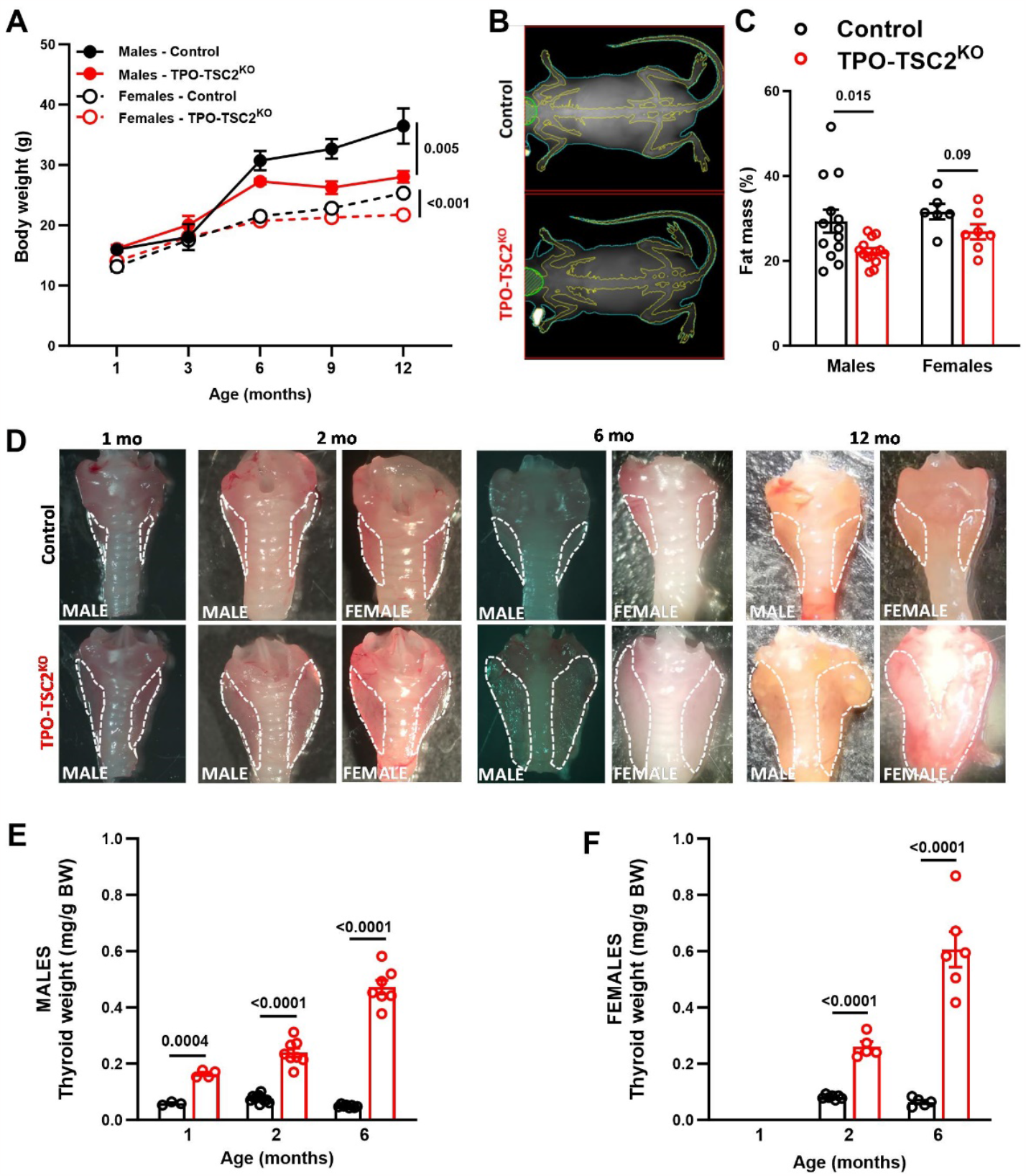
mTORC1 activation in thyrocytes induces a striking and progressive enlargement of the thyroid gland. **(a)** Body weight of Control and *TPO-TSC2^KO^* male and female mice; **(b)** Representative images of body composition assessment by Dual-energy X-ray (DEXA) of 12-month-old male mice; **(c)** DEXA fat mass quantification of 12-month-old male and female mice; **(d)** Representative images of thyroid glands of control and *TPO-TSC2^KO^* mice at 1, 2, 6 and 12 months of age; Thyroid weight normalized by body weight in **(e)** male and **(f)** mice. N = 5-10 mice per group. Data are shown as mean ± SEM. P values are shown in the figures determined by two-way ANOVA followed by Tukey’s posthoc test in **a** and unpaired two-tailed Student’s T-test in **c, e,** and **f**.

**mTORC1 increases *thyrocyte hypertrophy and proliferation***

We next assessed the thyroid gland structure and thyrocyte proliferation to understand how mTORC1 promotes thyroid growth. We measured ~100 follicles of female mice and observed a 3-fold increase in the follicular area (Fig 3A and B). This was explained by a 2-fold increase in both colloid and epithelium (thyrocyte) areas (Fig 3C and D). These effects were also noticeable using the individual median values (Fig 3E and G). Indeed, follicle area frequency distribution analysis revealed that *TPO-TSC2^KO^* mice have fewer small follicles (<1000 µm^2^) but a higher number of follicles with areas between 3000 and 6000 µm^2^ (Fig 3I). However, the epithelium-to-colloid ratio did not change (Fig 3H). Besides increasing in growth, mTORC1 enhanced thyrocyte proliferation assessed by Ki67 staining (Fig 4). The number of proliferating thyrocytes (TTF1^+^Ki67^+^ cells/TTF1^+^ cells) was 10-fold increase in female *TPO-TSC2^KO^* mice compared to controls at 6 months of age (Fig 4A and B). Because cyclin D3 is critical for thyrocyte proliferation [54, 81-83], we assessed its protein content and found that cyclin D3, but not D2, is increased in both males and females *TPO-TSC2^KO^* mice (Fig 4C and D).

**Figure 3.**
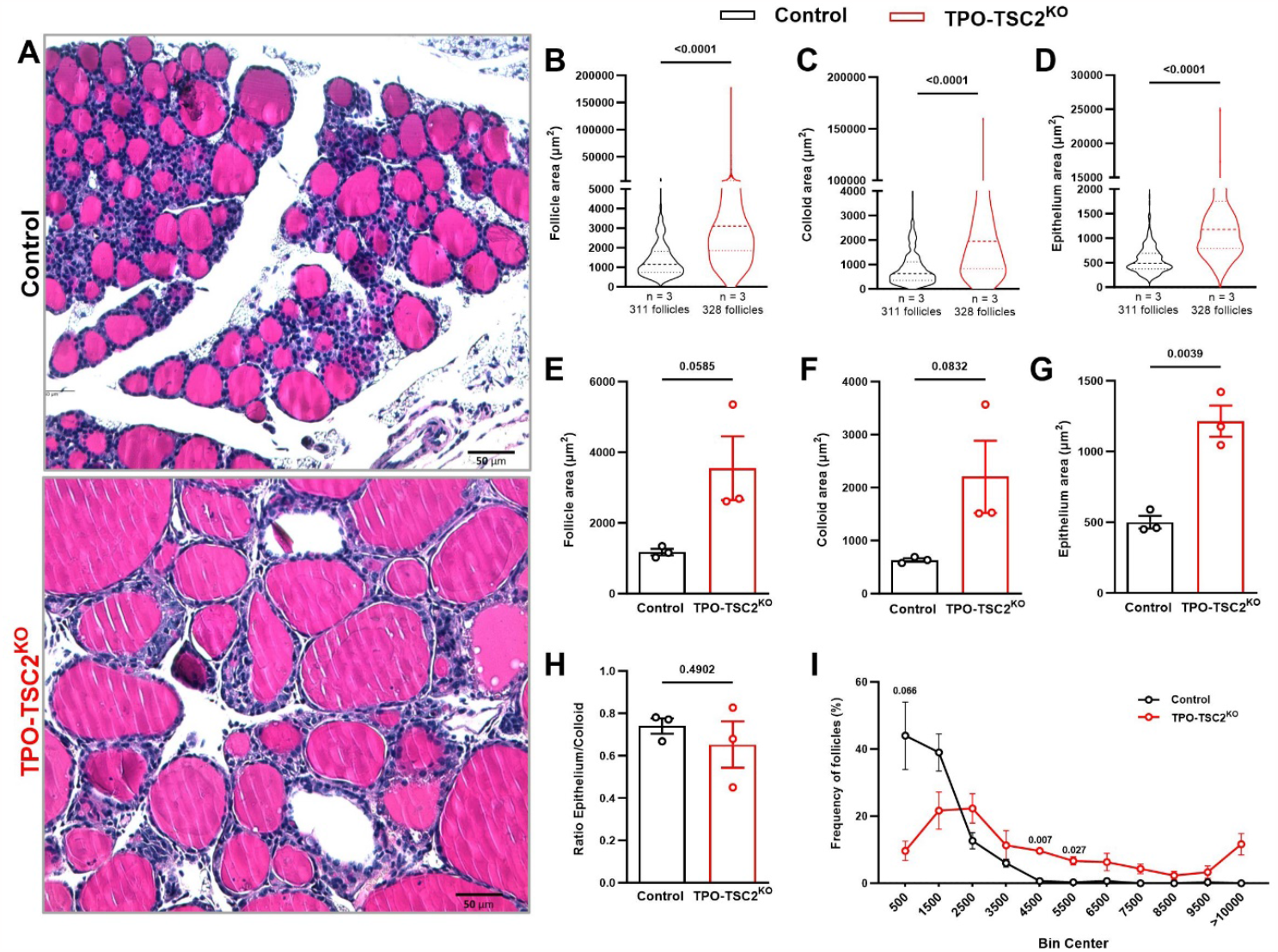
mTORC1 activation in thyrocytes leads to follicle enlargement. **(a)** PAS staining representative images of thyroid gland histological sections of control and *TPO-TSC2^KO^* mice. Scale bar: 50 µm; Quantification of **(b)** follicular area, **(c)** colloid area, and **(d)** epithelium area of all thyroid follicles (311 control and 328 *TPO-TSC2^KO^* follicles of 3 mice in each group). Median of **(e)** follicle area, **(f)** colloid area, and **(g)** epithelium area. Each plot represents the median of one mouse; **(h)** Epithelium/colloid area ratio and **(i)** Follicle frequency distribution according to its area. In **b, c** and **d** data are shown as a violin plot (median and Quartiles) and P values were determined by unpaired Mann-Whitney test. In **e, f** and **g** data are shown as mean ± SEM and p values determined by Student’s T-test. N = 3 mice per group.

**Figure 4.**
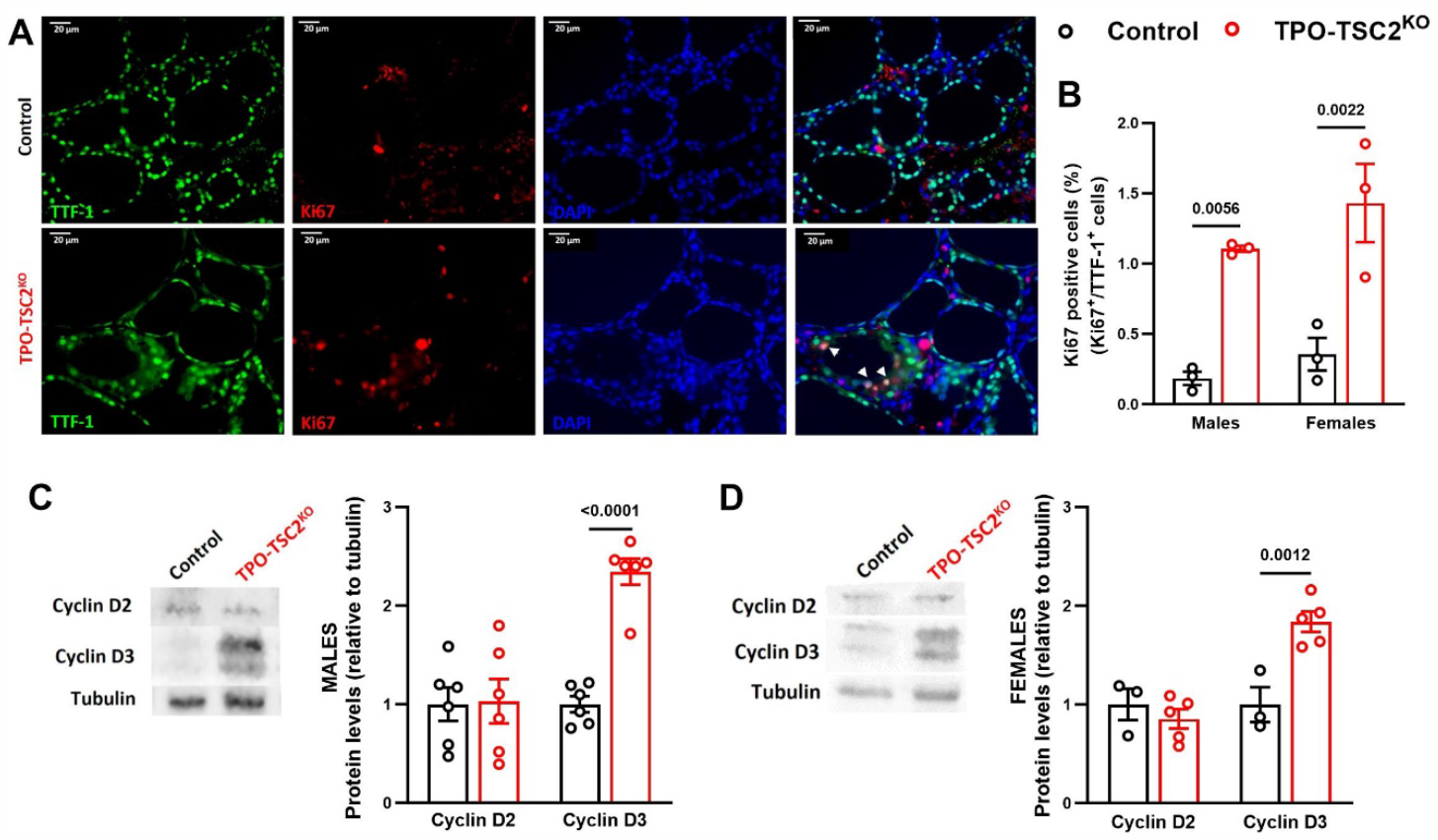
Intense thyrocyte proliferation in *TPO-TSC2^KO^* mice. **(a)** Representative images of TTF-1 (green), Ki67 (red) and DAPI (blue) immunostaining in thyroids of control and *TPO-TSC2^KO^* mice at 6 months of age. Scale bar: 20 µM. Arrows indicate proliferating thyrocytes (Ki67^+^/TTF-1^+^ cells); **(b)** Quantification of Ki67^+^/TTF-1^+^ cells in **a**. N = 3 mice per group; Cyclin D2 and Cyclin D3 protein expression in thyroids of 6 months old **(c)** males and **(d)** female mice. N = 3–6 mice per group. Data are shown as mean ± SEM. P values are indicated when less than 0.05, determined by unpaired two-tail Student’s T-test.

### mTORC1 regulates the expression of key regulatory genes in the thyroid gland

We next tested whether mTORC1 activation impacts gene expression in the thyroid of the *TPO-TSC2^KO^* mice. As expected, thyroid gland TSC2 mRNA levels are 60-90% reduced in *TPO-TSC2^KO^* mice of all ages and genders (Fig 5). Notably, NIS and TG mRNA levels are lower at 2, 6 and 12 months of age in male and female *TPO-TSC2^KO^* mice (Fig 5A-F). In addition, activation of mTORC1 promoted a remarkable decrease in NIS protein levels in 6-mo males and females (Fig 5G-H). In males, TPO expression tended to decrease at 2 and 6 months and was significantly decreased at 1-year old. TPO was less expressed in female *TSC2^KO^* mice at all ages (Fig 5A-F). While mTORC1 activation increased TSH receptor gene expression only in *TPO-TSC2^KO^* males (Fig 5A, C and E), it increased the type 1 deiodinase (DIO1) in females at 6 months (Fig 5D). To evaluate if this modified gene expression profile was a function of mTORC1-mediated alteration of thyrocyte differentiation, we also measured the expression of the two main transcription factors that determine thyrocyte identity and maturation, PAX8 and TTF1 [84]. We did not observe any difference in the mRNA levels of these genes in males or females *TPO-TSC2^KO^* mice at any age (Fig 5).

**Figure 5.**
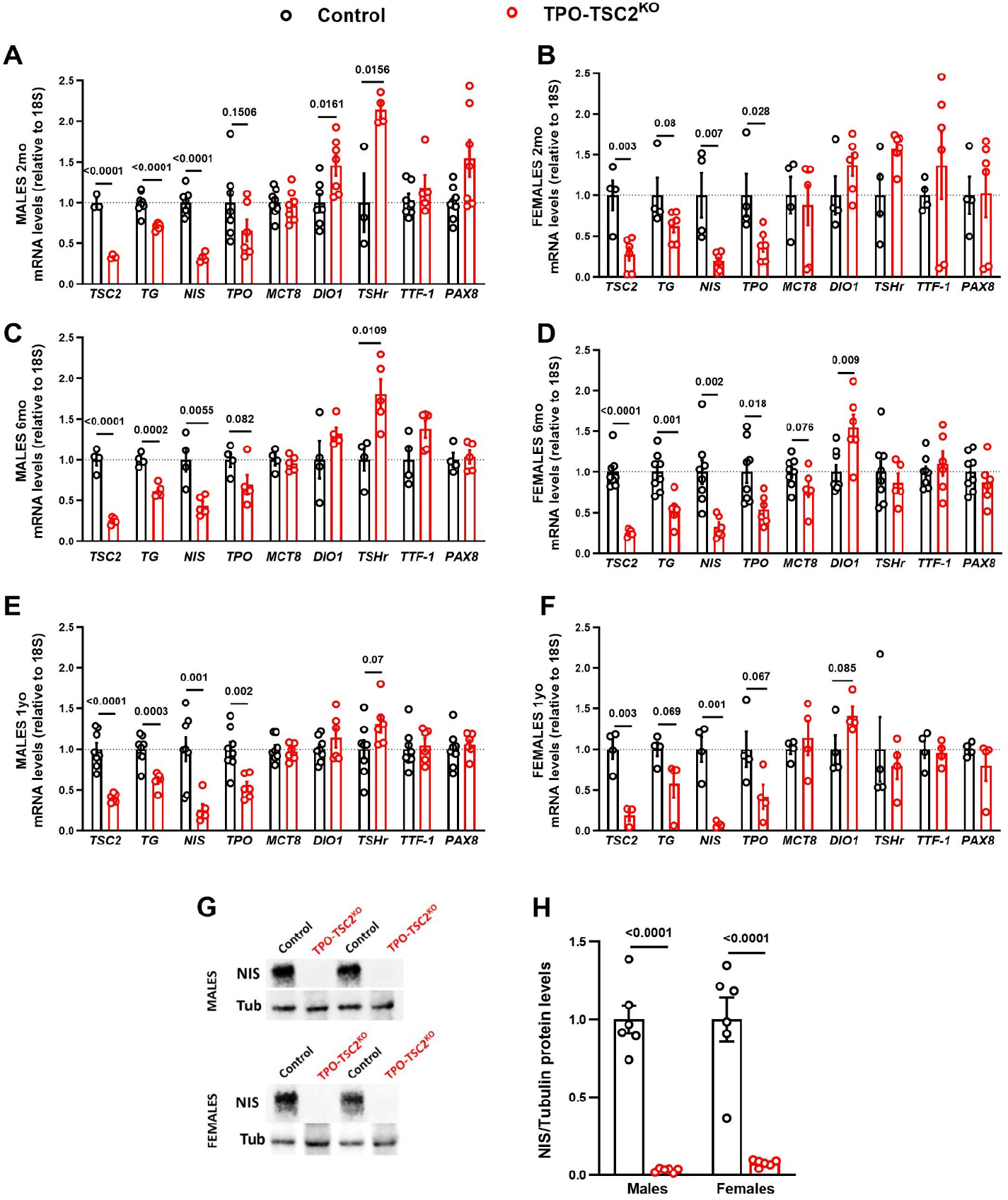
mTORC1 suppresses the expression of essential genes involved in thyroid hormone synthesis. Thyroid mRNA levels of 2-month-old **(a)** male and **(b)** female mice; Thyroid mRNA levels of 6-month-old **(c)** and **(d)** female mice; Thyroid mRNA levels of 1-year-old **(e)** male and **(f)** female mice; **(g)** Representative western blot images of NIS protein expression in males and females at 6 months of age and **(h)** quantification of images in **g**. N = 3–8 mice per group. Data are shown as mean ± SEM. P values are indicated above the bars and were determined by unpaired two-tailed Student’s T-test.

### Prolonged mTORC1 activation impairs thyroid function

Although mTORC1 activation altered the expression of key genes involved in TH synthesis, total T_4_, T_3_, and TSH plasma levels were normal in 2-month-old male *TPO-TSC2^KO^* mice (Table 3). At 6 months of age, both male and female *TPO-TSC2^KO^* mice had higher TSH levels with a trend to lower levels of T_4_ (p < 0.07 and p < 0.06, respectively; Tables 3 and 4). *TPO-TSC2^KO^* mice aged one year exhibited higher TSH and reduced T_4_ levels in both genders (Tables 3 and 4). In contrast, circulating T_3_ levels in *TPO-TSC2^KO^* mice were similar to controls throughout life.

**TABLE 3.**
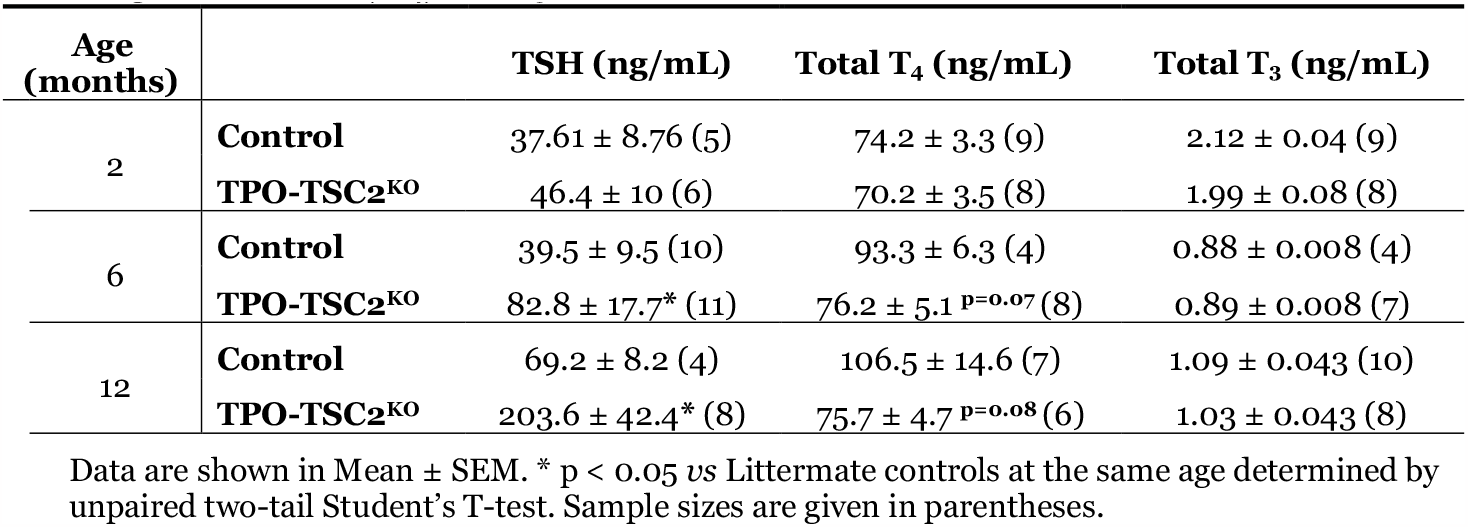
Plasma TSH, T_4_, and T_3_ in Control and TPO-TSC2^KO^ male mice.

| Age (months) |  | TSH (ng/mL) | Total T<sub>4</sub> (ng/mL) | Total T<sub>3</sub> (ng/mL) |
| --- | --- | --- | --- | --- |
| 2 | **Control** | 37.61 ± 8.76 (5) | 74.2 ± 3.3 (9) | 2.12 ± 0.04 (9) |
|  | **TPO-TSC2<sup>KO</sup>** | 46.4 ± 10 (6) | 70.2 ± 3.5 (8) | 1.99 ± 0.08 (8) |
| 6 | **Control** | 39.5 ± 9.5 (10) | 93.3 ± 6.3 (4) | 0.88 ± 0.008 (4) |
| | **TPO-TSC2<sup>KO</sup>** | 82.8 ± 17.7* (11) | 76.2 ± 5.1 $p=0.07$ (8) | 0.89 ± 0.008 (7) |
| 12 | **Control** | 69.2 ± 8.2 (4) | 106.5 ± 14.6 (7) | 1.09 ± 0.043 (10) |
| | **TPO-TSC2<sup>KO</sup>** | 203.6 ± 42.4* (8) | 75.7 ± 4.7 $p=0.08$ (6) | 1.03 ± 0.043 (8) |

Data are shown in Mean ± SEM. \* $p < 0.05$ vs Littermate controls at the same age determined by unpaired two-tail Student's T-test. Sample sizes are given in parentheses.

**TABLE 4.** Plasma TSH, T_4_, and T_3_ in Control and TPO-TSC2^KO^ female mice.

| Age (months) |  | TSH (ng/mL) | Total T<sub>4</sub> (ng/mL) | Total T<sub>3</sub> (ng/mL) |
| --- | --- | --- | --- | --- |
| 6 | **Control** | 32.2 ± 7.13 (10) | 118.7 ± 7.7 (6) | 0.86 ± 0.013 (5) |
| | **TPO-TSC2<sup>KO</sup>** | 71.1 ± 12.5* (8) | 100.2 ± 4.9 $p=0.06$ (7) | 0.87 ± 0.005 (6) |
| 12 | **Control** | 36.4 ± 10.2 (5) | 140.6 ± 14.1 (4) | 0.94 ± 0.008 (4) |
|  | **TPO-TSC2<sup>KO</sup>** | 160.8 ± 13.9*** (6) | 111.9 ± 6.4* (4) | 0.94 ± 0.007 (4) |

Data are shown in Mean ± SEM. \* $p < 0.05$ and \*\*\* $p < 0.0001$ vs Littermate controls at the same age determined by unpaired two-tail Student's T-test. Sample sizes are given in parentheses.

### Pharmacological Inhibition of mTORC1 reverses the TPO-TSC2^KO^ phenotype

To further demonstrate that the *TPO-TSC2^KO^* mice phenotype was due to enhanced mTORC1 activity in the thyrocytes, we pharmacologically inhibited mTORC1 with daily injections of Rapamycin, a well-known inhibitor of mTORC1. Both control and *TPO-TSC2^KO^* mice received Rapamycin (2 mg/kg *ip*) for 5 weeks from weaning (3 weeks old) until 2 months of age (Fig 6A). Rapamycin treatment successfully blocked mTORC1 activity (Fig 6D) and fully blunted mTORC1-mediated thyroid mass growth observed in the *TPO-TSC2^KO^* mice (Fig 6B and C). At the molecular level, Rapamycin administration reversed the changes in the expression of TG, TPO, NIS, Dio1, and TSHr (Fig 6E and F vs 5A and B). Total T_3_, T_4_, and TSH levels were not different between control and *TPO-TSC2^KO^* mice at 2 months of age (Table 3). However, we detected a decrease in blood T_4_ induced by Rapamycin treatment in the Control group (Vehicle 74.23 ± 3.26 *vs* 58.55 ± 2.77 ng/ml, p = 0.013), suggesting impaired thyroid function. These results suggest that the increase in thyroid growth and thyrocyte proliferation observed in *TPO-TSC2^KO^* mice occurs independently of TSH.

**Figure 6.**
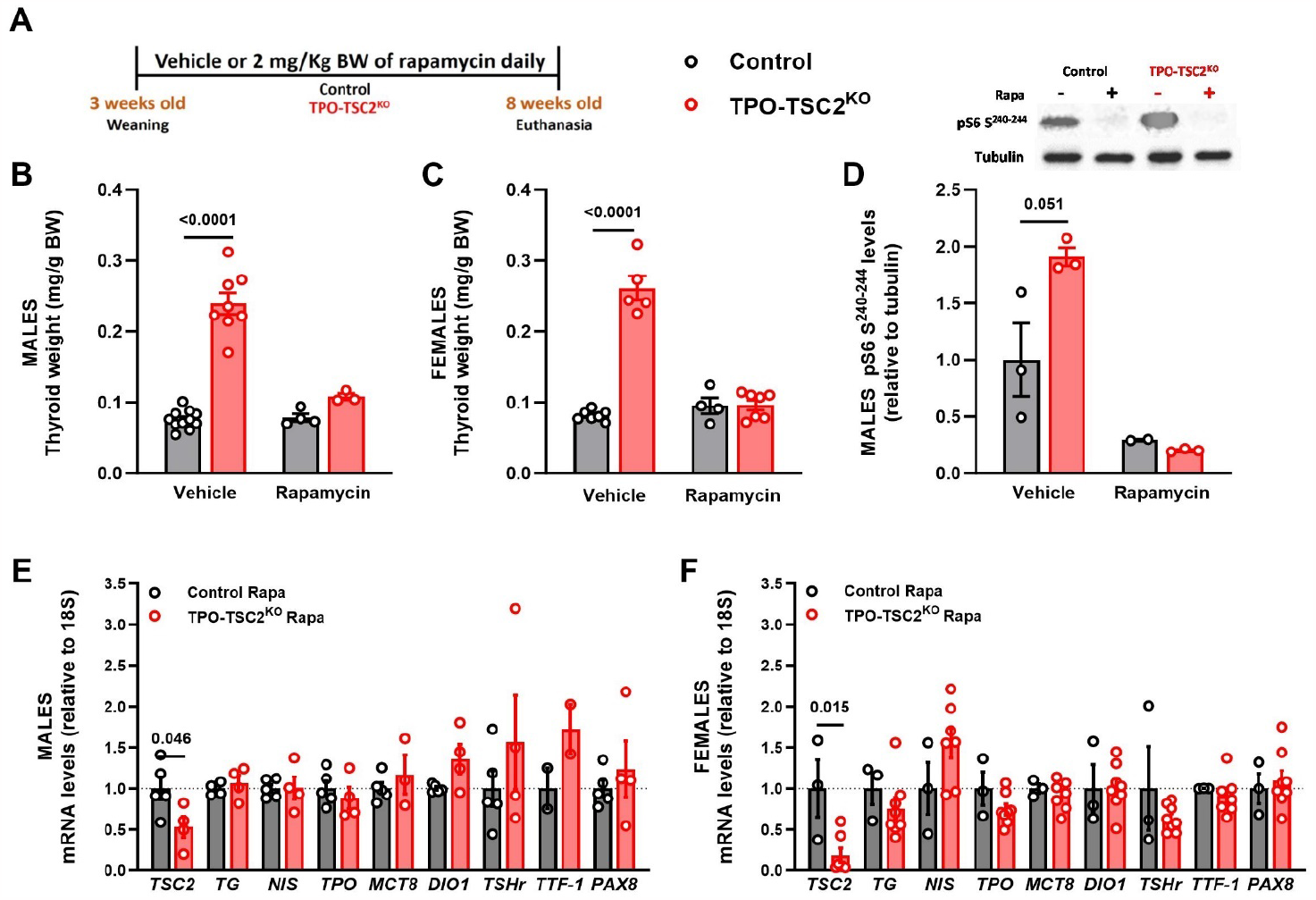
Rapamycin treatment blunts thyroid mass expansion and restores gene expression in the *TPO-TSC2^KO^*. **(a)** *TPO-TSC2^KO^* male mice were treated daily with 2 mg/Kg b.w. Rapa from 3 to 8 weeks of age; **(b)** Males and **(c)** Females thyroid weight normalized by body weight; **(d)** Western blotting of the thyroid gland for pS6 Ser^240-244^ (mTORC1 activity) and Thyroid mRNA levels in males **(e)** and females **(f)**. N = 3–6 mice per group. Data are shown as mean ± SEM. P values are indicated when less than 0.05, determined by unpaired two-tail Student’s T-test.

### TPO-TSC2^KO^ mice present aberrant thyroid follicles, inflammation, and tumors

Overactivation of mTORC1 led to the development of abnormal thyroids, characterized by multiple layers of thyrocytes and colloid between them (Fig 7A-C). Microscopic examination of thyroids from TPO-TSC2 knockout mice revealed nodular thyroid follicular disease with extensive cystic changes. Most of the follicles were lined by flattened or round oval follicular cells. Rare follicles exhibited focal centripetal papillary hyperplasia, and there were small areas with follicle breakdown with acute inflammation and histiocytic type reaction. After 12 months of age, mice with mTORC1 overactivation in thyroid developed tumors (Fig 7D-E).

**Figure 7.**
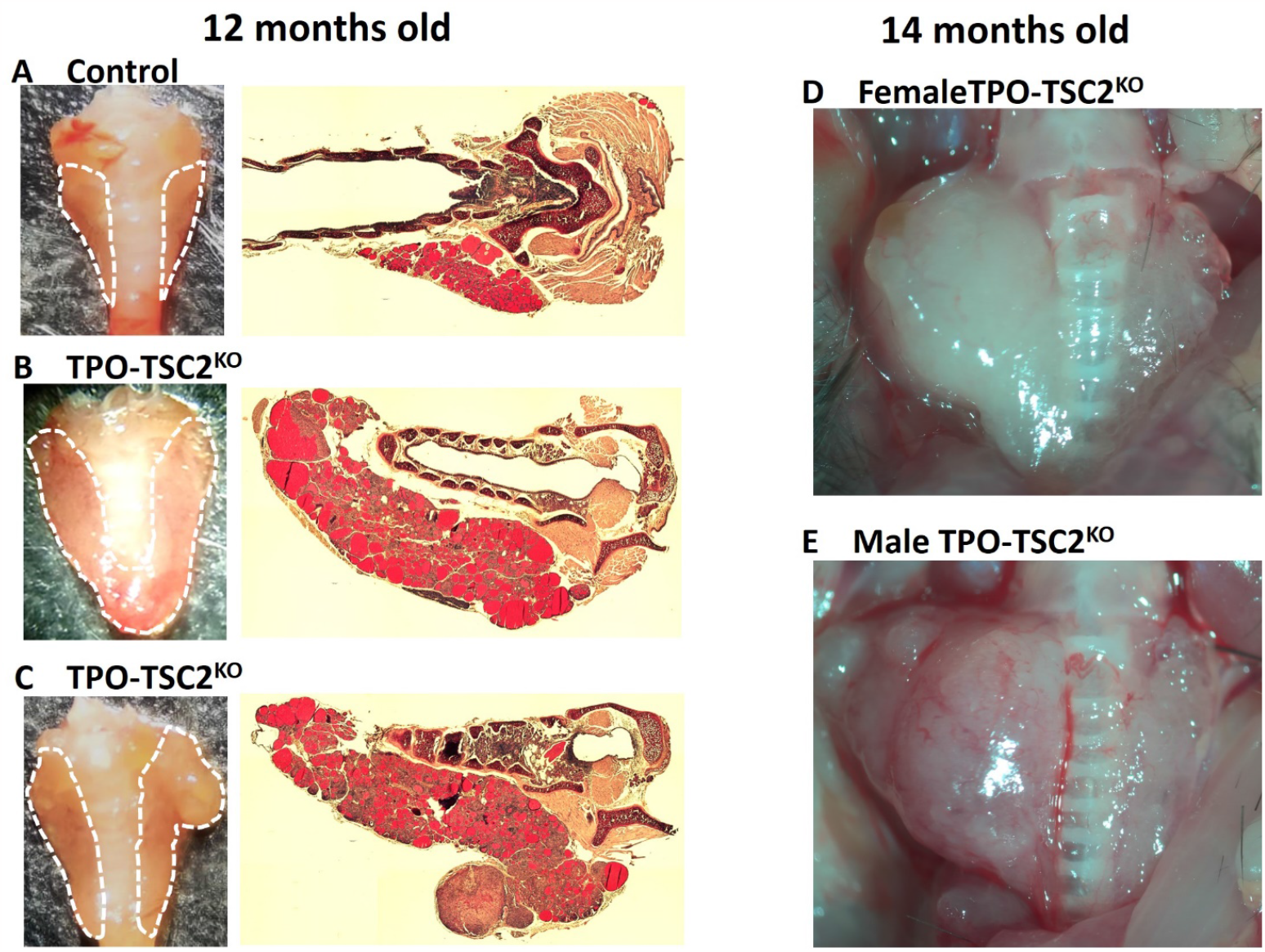
Twelve-month-old or older *TPO-TSC2^KO^* mice present aberrant thyroids. (a) Photo and photomicrographs of thyroid gland histological sections of 12 months old Control mice. (b) Photo and photomicrographs of thyroid gland histological sections of 12 months old *TPO-TSC2^KO^* mice. (c) Photo and photomicrographs of thyroid gland histological sections of 12 months old *TPO-TSC2^KO^* mice with a tumor. PAS staining. (d) Thyroid tumor in a 14 months old *TPO-TSC2^KO^* female. (e) Thyroid tumor in a 14 months old *TPO-TSC2^KO^* male.

## DISCUSSION

We demonstrate herein that activation of mTORC1 specifically in thyrocytes leads to a striking increase in thyroid gland weight, characterized by larger follicles filled with more colloid and thicker epithelium. In addition, mTORC1 drives intense thyrocyte proliferation. We also document that mTORC1 dramatically suppresses the expression of important genes for hormonogenesis such as *NIS, TPO*, and *TG*, impacting TH levels. Rapamycin treatment of *TPO-TSC2^KO^* mice blunted mTORC1-induced increase in thyroid growth and restored gene expression. This is the first *in vivo* study investigating the direct role of mTORC1 signaling activation specifically in thyrocytes.

The lack of *in vivo* models for investigating mTORC1’s role in thyrocyte physiology outside the context of thyroid cancer led to the generation of *TPO-TSC2^KO^* mice. It is well-established that mTORC1, a sensor of nutrients and growth factors, plays a crucial role in promoting cell growth and proliferation [7, 8]. Consequently, it is not surprising that *TPO-TSC2^KO^* mice exhibit thyrocyte growth and proliferation. Importantly, this mechanism operates independently of TSH, as TSH levels remain normal in young mice (less than 6 months of age). Previous *in vitro* and *in vivo* studies examining the effects of INS and IGF1 on thyrocytes indirectly suggested that mTORC1 regulates thyrocyte growth and proliferation. This inference arises from the fact that both INS and IGF1 activate the canonical PI3K-Akt-mTORC1 pathway. Research conducted in the 1980s and 90s, utilizing cell lines and primary thyrocytes from various animal species, demonstrated the cooperative action of IGF1/INS and TSH in promoting thyrocyte mitogenic activity [66-74]. At that time, the prevailing consensus was that IGF1/INS and TSH activate distinct intracellular pathways [36, 37]. However, more recent studies have reported that TSH can independently stimulate mTORC1, suggesting mTORC1 as a converging intracellular hub mediating thyrocyte responses to hormones and growth factors [32, 33, 75]. This newer evidence challenges the earlier notion of completely separate pathways, indicating a more integrated and complex signaling network governing thyrocyte behavior. The *TPO-TSC2^KO^* mouse model provides a valuable tool for dissecting the roles of mTORC1.

Mice overexpressing IGF1 and/or its receptor (IGF1R) in thyrocytes has been shown to induce thyroid gland growth and reduce the dependence on TSH [76]. This is substantiated by findings in mice with IGF1R deletion in thyrocytes (*IGF1R^KO^*), which exhibited impaired responses to TSH and goitrogenic diet [38]. Notably, mTORC1 activation was attenuated in the *IGF1R^KO^* mice, suggesting the involvement of mTORC1 in the adaptive responses to experimental goiter. However, a distinct phenotype was observed in double knockout mice for INS and IGF1 receptors (*IRKO/IGF1R^KO^*) [40]. Despite having smaller thyroids in the neonatal period, by postnatal day 14, these mice paradoxically displayed increased thyrocyte proliferation. The authors hypothesized that elevated circulating TSH levels compensated for the loss of IGF1 and INS signaling, inducing thyrocyte hyperplasia [40]. An independent study on *IGF1R^KO^* mice reported a similar TSH compensation phenomenon [39]. Our data from the *TPO-TSC2^KO^* mice align with the notion that mTORC1 is essential for thyroid proliferation and growth [38] but also indicate that TSH-induced compensation might depend on thyrocyte mTORC1 activity [39, 40]. Furthermore, mTORC1 appears to mediate thyrocyte proliferation in adaptive responses to iodine deficiency, as chronic *in vivo* treatment of mice with the rapalog RAD001 (Everolimus) during goitrogenic conditions abolished the hyperplastic thyrocyte response [32]. However, it’s noteworthy that the authors did not observe any effects of rapamycin treatment on thyroid gland growth. The proposed uncoupling of mTORC1 action on proliferation and growth, as suggested by the authors, is not supported by our *TPO-TSC2^KO^* mouse model.

*In vivo* data demonstrating the impact of the PI3K-Akt-mTOR pathway on TH synthesis and secretion is currently limited and subject to controversy [31, 42-45]. This controversy can be attributed to the constraints of *in vitro* conditions, which fail to replicate the organized thyrocyte structures as follicles and consequently do not fully represent thyroid gland physiology [26, 77]. Over the last two decades, research has primarily focused on understanding the regulation of NIS expression and activity by mTORC1 and/or mTORC2, particularly in the context of more aggressive thyroid cancers where downregulation of NIS renders them refractory to radioiodine therapy [46, 78, 79]. In contrast to the observed synergism and cooperation between TSH and INS/IGF1 in thyrocyte proliferation and growth, initial studies using FRTL-5 and PCCL3 cell lines suggested opposing actions of these hormones in the regulation of NIS [31, 44, 45]. However, recent investigations using primary human thyrocytes have demonstrated additive effects of TSH and IGF1 on the mRNA expression of TG, TPO, Dio2 and NIS [42, 43]. Our present data, derived from the *TPO-TSC2^KO^* mice, indicate that mTORC1 downregulates NIS gene and protein expression. This finding aligns with observations that high doses of Everolimus (four oral doses of 5 mg/kg body weight in 24 h) increased thyroid iodide uptake by 50% in Wistar rats [44]. However, it is important to note that such high doses may also lead to the blocking of mTORC2 activity.

In the thyroids of *TPO-TSC2^KO^* mice, besides *NIS*, the downregulation of other genes positively regulated by TSH, such as *TPO* and *TG*, was evident. This mTORC1-mediated effect was observed at both normal TSH levels (1 and 2 months of age) and elevated TSH levels in the blood (6 and 12 months-old mice). TSH is known to activate signal transduction pathways that intricately regulate the transcription and activity of Thyroid Transcription Factors (TTFs), including TTF1, TTF2, and PAX8. These factors collectively control the gene expression of various thyroid-specific functional proteins, including *NIS, TG, TPO, DIO1, and DIO2* [80-82]. TTFs typically converge on the cis regulatory elements of these target genes and regulate their expression in a feedforward fashion [81]. Notably, in primary human thyrocytes, knockdown of *TTF1, TTF2*, and *PAX8* genes has been shown to inhibit TSH-stimulated *NIS, TG, TPO,* and *DIO2* mRNA expression [83]. However, we did not observe a decrease in the expression of *TTF1* or *PAX*-8, suggesting that mTORC1 effect on *NIS, TG* and *TPO* are not TTFs-mediated.

The development of the thyroid gland occurs rapidly during the last third of gestation, and its maturation continues until the second postnatal week [65]. In our model, mTORC1 activation is expected to begin when TPO is expressed during embryonic development (E14.5), subsequent to TTF1 and PAX8 expression [59]. We did not observe any noticeable difference in TTF1 (Fig 5) and PAX8 (not shown) nuclear protein staining in the *TPO-TSC2^KO^* mice. This suggests that mTORC1 is likely to directly modulate the expression of NIS, TPO, and TG rather than inducing dedifferentiation of thyrocytes. However, it’s conceivable to argue that the loss of expression of these critical genes may indicate a partial loss of thyrocyte identity, despite the maintenance of follicle structure until the mice age to one year. In contrast and intriguingly, *TSHR* and *Dio1* were upregulated, even in the presence of normal TSH levels, possibly as a compensatory mechanism to counteract the impaired synthesis of TH.

It is noteworthy that T4 and T3 plasma levels are normal in the *TPO-TSC2^KO^* mice at 1 and 2 months of age despite the downregulation of *NIS, TPO* and *TG* expression. Only when animals are 6 months or older, T4 slightly decreases and TSH increases in male and female mice. It is conceivable to speculate that the striking increase in thyrocyte proliferation and mass (thyroid epithelium layer), and increases in follicle content compensates for the decrease in TH synthesis. Indeed, TH plasma concentration per gram of thyroid (T4/thyroid weight ratio) is dramatically decreased in the *TPO-TSC2^KO^* mice compared to littermate controls.

## CONCLUSION

mTORC1 not only plays a pivotal role in controlling thyrocyte proliferation but also significantly influences thyrocyte function. Based on the results presented, we conclude that the activation of mTORC1 pathway can decrease the expression of NIS, TPO and TG, ultimately to reduced T_4_ secretion. Future research will be directed towards identifying the specific intracellular targets of mTORC1 that mediate downregulation of TH synthesis genes, and investigating whether mTORC1 is involved in TH secretion.

